# Chloroform release from ageing cells and *Drosophila* DJ-1 mutants

**DOI:** 10.1101/2025.01.13.632713

**Authors:** Theo Issitt, Timothy Johnston, Chris Ugbode, Juste Grumulaityte, Amy Harmens, William J Brackenbury, Sean T Sweeney, Kelly R Redeker

## Abstract

Volatile Organic Compounds (VOCs) offer potential for non-invasive diagnosis as biomarkers of disease and metabolism. In complex biological matrices, such as breath however, identifying useful biomarkers from hundreds, or even thousands of VOCs can be challenging. Models of disease, such as cellular or animal models, offer a means to elucidate VOC metabolisms, for accurate targeted studies in patient samples. Neurodegenerative conditions, such as parkinsons have been associated with changed VOCs, offering a potential for early diagnostics and interventions improving treatments and outcomes for patients. Here, three separate models including; human HEK-293t cells, isolated primary rat glial cells, *Drosophila* flies (wild type and a mutant of the parkinson’s associated gene, DJ-1) were grown for an extended period and levels of the VOC chloroform (CHCl_3_) investigated using custom static headspace sampling chambers. Samples were analysed using targeted gas chromatography mass spectroscopy over time to generate metabolic flux values and chloroform shown to dramatically increase in all models as they aged. HEK-293t cells revealed a 60-fold increase after 10 weeks, glial cells revealed a 10-fold increase after 3 to 4 weeks and DJ-1 mutant flies revealed significant increases compared to control flies at 4 weeks. These results, taken together, indicate that chloroform release is related to ageing in these models and may provide a target for neurodegenerative studies moving forward. We present here the first evidence of chloroform being actively produced by human and rat cells and the first observation of volatile metabolisms in *Drosophila*.

## Introduction

Volatile Organic Compounds (VOCs) diffuse readily through cellular membranes and liquids and are released and consumed through cellular metabolism, offering potential non-invasive diagnosis [1,2]. VOCs define odour and examples of smell being used for the diagnosis of neurodegenerative conditions have encouraged efforts for non-invasive diagnostic technologies [3,4]. Despite significant advances in understanding of VOC mechanisms [5,6] and technologies for diagnosis via VOCs [7], applications within the clinic are still limited [1,3].

Neurodegenerative diseases, such as parkinsons, are defined by a change in function of the neurological system. There are many causes of neurodegenerative disease [8] but early intervention can improve outcomes and therapeutic options for patients [8]. Because the neurological system is highly metabolically active [9], VOCs, as gaseous metabolites therefore offer an opportunity for non-invasive diagnosis.

A critical need to accelerate translation of this tool to the clinic is identification of biomarkers, linked to neurodegenerative conditions, which would allow targeted methodologies to simplify analysis of complex biological matrices, such as breath or blood. VOCs from cells in culture have been shown to distinguish between tissue of origin and pathology [10], cell status [11], or response to stress [12,13]. A number of VOC studies from the breath of animals have shown VOCs can diagnose a range of conditions, such as infection in mice [14] or pigs [15] and gastrointestinal and hepatic disease in rats [16]. Cellular and animal models of disease are therefore useful tools for potential biomarker identification for validation in human studies, framing the purpose of this work.

In this study we investigated the release of chloroform (CHCl_3_) in the headspace of aging cells and flies as a result of a chance observation when profiling VOCs from HEK293t cells which were passaged for an extended period of time. We hypothesised that cells accumulate cellular products over time and went about to test if chloroform release in aging cells was observed in terminally-differentiated cell culture sourced from rat brain. Furthermore, we aimed to establish if this could be observed in a model of neurological dysfunction, such as parkinsons, using *Drosophila melanogaster*. In parkinsons disease, and a range of neuronlogical conditions the clearing of metabolic products such as ROS is inhibited [17]. Protein degylcase (DJ-1) also known as PARK7 has been used as a model of neurological dysfunction in drosophila as it is altered in parkinson’s disease [18]. DJ-1 protects against accumulation of reactive oxygen species (ROS), and DJ-1 mutant flies have increased levels of ROS [19] and here we hypothesize that DJ-1 may present with elevated levels of chloroform over time in comparison to controls.

## Methods

### Cell Culture and cellular isolation

HEK-293t were grown in Dulbecco’s Modified Eagle Medium (DMEM, Thermo Scientific, Waltham, MA, USA), 25 mM glucose, supplemented with L-glutamine (4 mM) and 5% foetal bovine serum (Thermo Scientific, Waltham, MA, USA). Cell were passaged twice a week.

To initiate the volatile collection for HEK293t, cells were trypsinised and ∼500,000 cells were seeded into 8 mL complete media. Cells were then allowed to attach for 3 h, washed with warm PBS 2× and an 8 mL treatment media was applied. Volatile headspace sampling was performed 24 h later as previously described [10].

### Glial cell isolation and maintenance

Timed-mated female Wistar rats (Charles River UK) (RRID:RGD_737929) were maintained in accordance with the UK Animals (Scientific Procedures) Act (1986). Cortices were dissected from postnatal day 1 (P1) mixed sex rat pups. Animals were euthanised using pentobarbital injection followed by cervical dislocation, according to Home Office guidelines. Cortical cell suspensions were obtained as previously described [20] and cytosine arabinoside (AraC, 2.4 μM final concentration) was added to the growth medium at 1 day in vitro (DIV) [21]. Glial cultures were confirmed following cells establishment in culture (20 days) and no neurons were observed in culture.

### Fly strains and husbandry

Wild-type and transgenic strains were reared and maintained on standard yeast–agar– cornmeal medium [22] and kept at 25°C for 12 hour light/dark cycles. Wild-type, white-eyed flies (*w*^*-118*^) and *DJ-1β*^*Δ93*^ mutants [23] were maintained for the indicated times prior to analysis, with the flies flipped onto fresh food 1-2 times per week.

### Static volatile headspace sampling

Detailed methods of static headspace sampling have been previously published [10,12]. Briefly, for cells in culture, 8ml fresh media was applied to cells in 10cm petri dishes 24 hours before sampling. Upon sampling initiation, 5ml media and petri dish lid was removed and dishes placed in static chambers. Both flies and cell cultures in headspace chambers were placed on a rocker on lowest setting with lab air flowing through the chamber for 10mins at 750ml per min. Chambers were then sealed and time point 0 samples taken, T1 samples were then taken 2 hours later for cells in culture and 4 hours for flies. 2 hours sampling point for flies was insufficient for observation of adequate signal, possibly due to passive respiration. All samples were collected through pressure differential into pre-evacuated, electropolished, stainless steel canisters.

### GC-MS, Calibration and Peak Analysis

Methods have been previously published [10,12]. Collected canister samples were transferred to a liquid nitrogen trap through pressure differential. Pressure change between beginning and end of “injection” was measured, allowing calculation of the moles of canister collected air injected. Sample in the trap was then transferred, via heated helium flow, to an Aglient/HP 5972 MSD system (Santa Clara, CA, United States) equipped with a PoraBond Q column (25 m × 0.32 mm × 0.5 μm film thickness) (Restek©, Bellefonte, PN, United States). Targeted samples were analyzed in selected ion monitoring (SIM) mode. The mass spectrometer was operated in electron impact ionization mode with 70 eV ionization energy, and transfer line, ion source, and quadrupole temperatures of 250, 280, and 280, respectively. A known standard was used with masses 83/85 for identification of chloroform. All samples were analysed within 6 days of collection. The oven program for both SIM were as follows: 35°C for 2 min, 10°C/min to 155°C, 1°C/min to 131°C, and 25°C/min to 250 with a 5 min 30 s hold.

### Statistics

Stastical analysis and figure arrangement was performed in Graphpad Prism (v10). Students T-test was performed for Figure 1A and B, one way ANOVA with bonferonni post host analysis was performed for Figure 1D.

**Figure 1.**
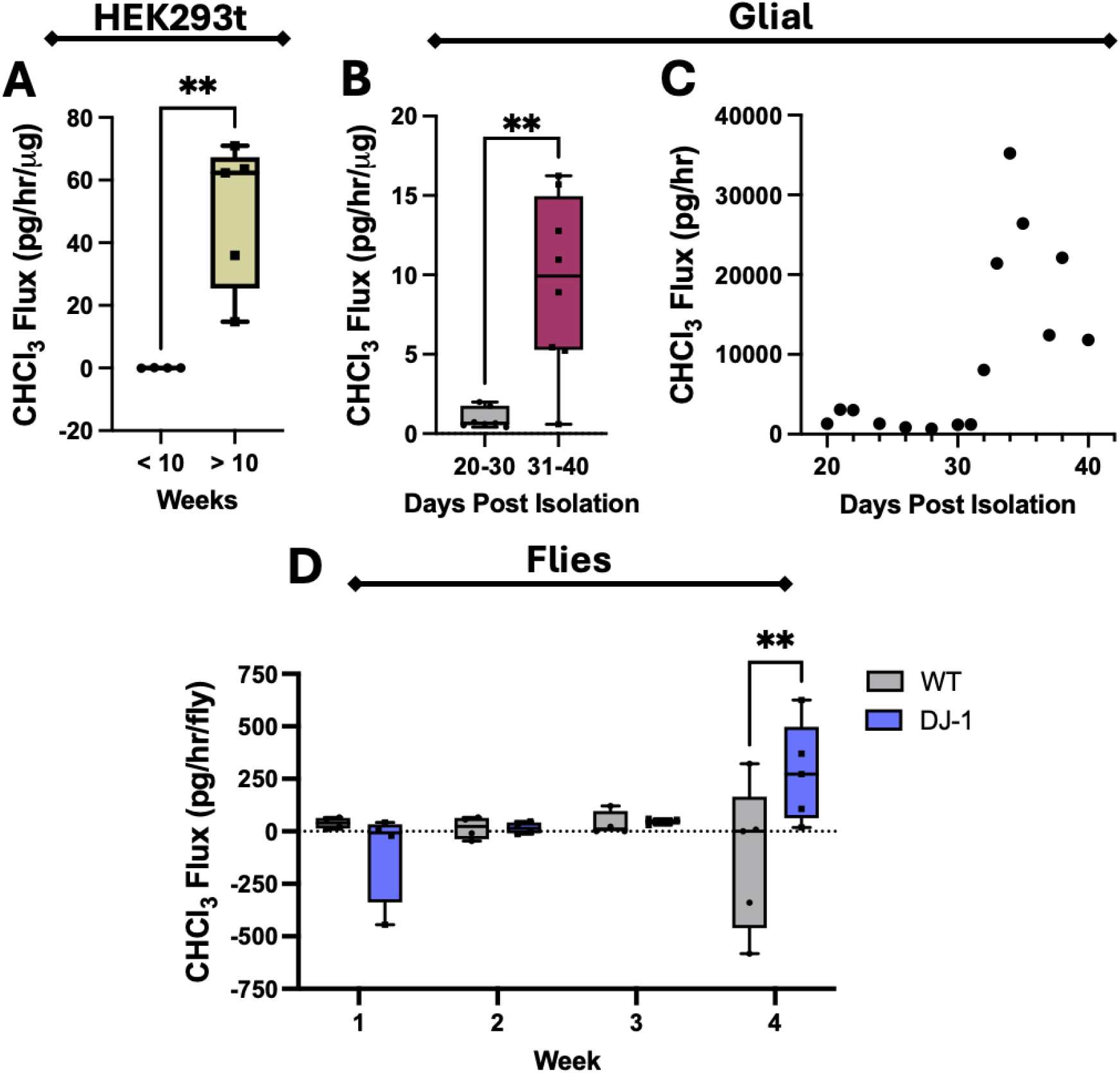
Chloroform (CHCl_3_) flux in aging HEK293t cells, rat glial cells and wild-type (WT) or DJ-1 mutant *Drosophila melanogaster* flies. **(A)** CHCl_3_ flux (pg CHCl_3_/hour/μg total protein), with media subracted for human HEK-293t cells below (<10, n =4) or above (>10, n = 5) passage 20 (passages on average twice a week). **(B)** CHCl_3_ flux (pg CHCl_3_/hour/μg total protein), media subtracted for primary rat glial cells for days post isolation(20-30, n = 7; 31-40, n = 8). **(C)** CHCl_3_ flux (pg CHCl_3_/hour) for glial cells days post isolation. **(D)** CHCl_3_flux(pgCHCl_3_/hour/per fly) of WT or DJ-1 mutant flies at 1,2, 3 (n = 4) and 4 (n = 5) weeks of age. Students T-test performed for **A** and **B**, two-way AN OVA with Bonferroni post-hoc analysis performed for **D**; ** p = 0.001.

## Results

HEK-293t cells passaged over 20 times (10 to 15 weeks in culture) produced chloroform at significantly higher levels than cells below this age (0.05±0.046 pg/hr/μg for <10 weeks and 49.5±23.5 pg/hr/μg for >10 weeks in culture).

Rat glial cell cultures were confirmed following isolation at day 20. Glial cells released chloroform with cells over 30 days in culture releasing significantly more than those between 20 and 30 days (Figure 1B, 0.95±0.64 pg/hr/μg for 20-30 days and 9.47±5.49 pg/hr/μg for 30-40 days). Significant release of chloroform in glial cells occurred at 33 days with slight increases occurring in cells between 30 and 32 days (Figure 1C).

Over 4 weeks, control flies showed no significant alterations in chloroform flux whereas the DJ-1 mutants revealed significant production of chloroform at week 4 compared to wild type flies (Figure 1C, -118.22 ± 349.12 pg/hr/μg vs 278.72 ± 237.39 pg/hr/μg).

## Discussion

We present here the first example of chloroform release from human cells, rat glial cells and the first documented example of volatile metabolisms from *Drosphillia melongasta*. We have previously investigated chloroform flux in HEK-293t and other cells in cultures, as well as the response of these cells to hypoxia and chemotherapeutic stress [10,12]. Modulation of chloroform metabolism was not observed from between cell lines or under conditions of stress/starvation as well as no active metabolism observed in the breath of mice [10].

The strikingly large amount of chloroform released by both HEK-293t and glial cells as they age (10-50x times greater flux) could be a result of accumulating chloride containing compounds, potentially as intracellular waste [17]. In culture, glial cells, which are highly metabolically active, accumulate waste which would normally be cleared by macrophages and immune cells and poor waste management is a hallmark of neurodegenerative conditions [24].

While, we could find no studies investigating chloride accumulation in our models over time and further studies are required, chloride accumulation is associated with brain disorders, trauma and other neurological conditions [25,26] and ionic dysregulation is a hallmark of metabolic dysfunction [27–29]. Furthermore, there is a lack of studies investigating chloride regulation in the brain but there is mounting evidence that disorders of the nervous system are caused by abnormal homeostasis of the intracellular concentration of Cl− [26,30].

We observed positive release of chloroform from DJ-1 mutant flies over 4 weeks, however few flies survived into week 5 and we were unable to measure past 4 weeks (over 28 days) which would have allowed clearer comparison with glial cells.

Few sources observe metabolic production of chloroform but it has been observed in fungi [31] and is persistent in the environment drinking water, driven by human pollution [32]. Cellular accumulation of chloride ions or chloride containing compounds in culture could be a source of chloroform. Carbon tetrachloride (CCl4) has been demonstrated to be a source of chloroform through cytochrome P450 dependent metabolism of carbon tetrachloride in hepatic microsomes [33], degradation in mouse liver [34] and aerobic transformation by poplar cells [35]. Neurons and glial cells are able to metabolise CCl4 [36] and this could be a potential source of the observed chloroform release, however, presence of CCl4 in our models requires further investigation.

In conclusion, we have demonstrated positive production of chloroform in ageing cells and DJ-1 mutant flies. Chloroform therefore presents as a potential biomarker to target in diseases associated with ageing, such as parkinson’s disease.

## Acknowledgements

The work was also supported by the White Rose Mechanistic Biology Doctoral Training Program, supported by the Biotechnology and Biological Science Research Council (BBSRC) BB/M011151/1. The authors would like to acknowledge the support provided by Mark Bentley in the University of York Department of Biology workshop.

